# Molecular basis and functional consequences of the interaction between the Base Excision Repair DNA glycosylase NEIL1 and RPA

**DOI:** 10.1101/2021.05.11.443618

**Authors:** Rémy A. Le Meur, Turner J. Pecen, Kateryna V. Le Meur, Zachary D. Nagel, Walter J. Chazin

**Affiliations:** Departments of Biochemistry and Chemistry, and Center for Structural Biology, Vanderbilt University, Nashville, TN 37240-7917, USA; Department of Environmental Health, Harvard T.H. Chan School of Public Health, Boston, MA, 02115-6009, USA; Biological NMR Technological Platform, Institut Pasteur, CNRS-UMR-3528, 75015 Paris, France

## Abstract

NEIL1 is a DNA glycosylase that recognizes and initiates base excision repair of oxidized bases. The ubiquitous ssDNA binding scaffolding protein replication protein A (RPA) modulates NEIL1 activity in a manner that depends on DNA structure. Interaction between NEIL1 and RPA has been reported, but the molecular basis of this interaction has yet to be investigated. Using a combination of NMR spectroscopy and isothermal titration calorimetry (ITC), we show that NEIL1 interacts with RPA through two contact points. An interaction with the RPA32C protein recruitment domain was mapped to a motif in the common interaction domain (CID) of NEIL1 and a dissociation constant (Kd) of 200 nM was measured. A substantially weaker secondary interaction with the tandem RPA70AB ssDNA binding domains was also mapped to the CID. Together these two contact points reveal NEIL1 has a high overall affinity (Kd ∼ 20 nM) for RPA. A homology model of the complex of RPA32C with the NEIL1 RPA binding motif in the CID was generated and used to design a set of mutations in NEIL1 to disrupt the interaction, which was confirmed by ITC. The mutant NEIL1 remains catalytically active against ionizing radiation-induced DNA lesions in duplex DNA *in vitro*. Testing the functional effect of disrupting the NEIL1-RPA interaction *in vivo* using a Fluorescence Multiplex-Host Cell Reactivation (FM-HCR) reporter assay revealed that RPA interaction is not required for NEIL1 activity against oxidative damage in duplex DNA, and furthermore revealed an unexpected role for NEIL1 in nucleotide excision repair. These findings are discussed in the context of the role of NEIL1 in replication-associated repair.

## Introduction

Reactive Oxygen Species (ROS) generated by endogenous and exogenous agents provide a constant source of damage to DNA (1). If left unrepaired, oxidation of bases caused by ROS can alter the base-pairing of DNA. Mispairing, in turn, can lead to fork instability and generation of mutations that can ultimately cause a variety of diseases including cancer, expedited ageing and neurodegeneration (2, 3). Single-strand DNA (ssDNA), which exists transiently during replication, transcription and recombination, is highly sensitive to ROS. The most common reactions are cytosine deamination into uracil, alkylation of adenine and cytosine, and spontaneous depurination and depyrimidination (1, 4–6). These and other oxidized bases are primarily repaired by the Base Excision Repair (BER) pathway. Although repair of oxidized bases are essential for genome stability, incomplete repair in the context of replication can lead to highly toxic strand breaks (7). Therefore, tight coordination between BER and the replication machinery is essential for genome maintenance in replicating cells.

NEIL1 is a DNA glycosylase that initiates BER of oxidized bases and is critical for pre-replicative DNA repair (8). Among the 5 mammalian DNA glycosylases identified to initiate BER of an oxidized base (OGG1, NTH1, NEIL1, NEIL2, NEIL3), only NEIL1 and NEIL2 are effective on both double stranded (ds) DNA and ssDNA (9, 10). NEIL1 expression is up-regulated in S-phase and interacts not only with BER enzymes, such as PolB, Lig3 and XRCC1, but also with replication enzymes such as PCNA and RPA (11). PCNA stimulates NEIL1 excision of damaged bases. This interaction suggests a role for NEIL1 in surveillance of DNA as the replication fork progresses (11). RPA stimulates NEIL1 activity when the damage is present in dsDNA near a ssDNA junction, as in pre-replicative DNA. Conversely, NEIL1 inhibits excision of a damaged base in ssDNA (12). This function is believed to help prevent the formation of toxic strand breaks. Thus, there is substantial evidence that the interaction between RPA and NEIL1 is crucial to the regulation of BER during replication; however, a detailed molecular understanding of this interaction and its consequences is not yet available.

The interaction between RPA and NEIL1 has been previously characterized using a combination of deletion mutagenesis, co-immunoprecipitation, in-vitro pull-down, far western and fluorescence binding assays (12). Binding to RPA was mapped to the RPA70 sub-unit and was shown to involve NEIL1 residues 289-349 within the Common Interaction Domain (CID). Here we report a detailed study of the molecular basis of NEIL1-RPA interaction using a combination of NMR, isothermal titration calorimetry (ITC), computational and cell-based Fluorescence Multiplex-Host Cell Reactivation (FM-HCR) reporter assays. The design, generation and validation of a specific NEIL1 mutant inhibiting the physical interaction between NEIL1 and RPA provided a valuable reagent to test the functional effects of suppressing this interaction. Leveraging the unique ability of the FM-HCR to simultaneously probe multiple DNA repair pathways, we discovered an unexpected role for NEIL1 in nucleotide excision repair (NER). These results suggest potential roles for NEIL1-RPA interaction in replication-associated base excision repair.

## Materials and Methods

### Plasmids

Cloning of RPA constructs is described elsewhere (13–16). NEIL1 expression vectors were a kind gift from Drs. Muralidnar Hedge and Sankar Mitra. The NEIL1 K319E/R323E/R326E/ R329E mutant was generated using the Quick Change® protocol as previously described (17). PCR primers were 5’-CGAGACACGAGAGGCAAAGGAAGACCTTCCTAAGAGGAC-3’ and 5’-CCTCTCGTGTCTCGGAAGGGGCCTCGCTTGGA-3’. Correct incorporation of the mutations was confirmed by DNA sequencing (Genewiz). The WT and mutant genes were then amplified from these expression plasmids using the PCR primers 5’-GCGCTAGCAGCACCCATATGCCTGAG-3’and 5’-ACGAAGCTTTCAGTGGTGGTGGTGGT-3’and subcloned into the pMaxCloning vector (Lonza, VDC-1040) via NheI and BamHI sites to generate the pMax_WT_NEIL1 and pMax_Mutant_NEIL1 expression plasmids that were utilized throughout this analysis.

### Expression, purification of NEIL1 and RPA

RPA70N, RPA32C, RPA70AB and full length RPA were expressed and purified as previously described (14, 15, 18, 19). Wild-type and mutant NEIL1 proteins were expressed from a pET22b vector harboring the wild-type or mutant NEIL1 gene fused with C-terminal 6-His tag transformed into Rosetta 2 (DE3) *E. coli* cell line. A fresh colony was used to inoculate a starter culture of 100 mL of autoclaved Terrific Broth (TB) medium supplemented with ampicillin in 250 mL baffled flask. After overnight growth at 37 °C with shaking at 230 rpm, 15 mL of starter culture was used to inoculate each 1L of TB culture medium supplemented with ampicillin in a 2L baffled flask. Growth was carried out at 37 °C with shaking at 230 rpm until OD_600_ reached 0.8. Temperature was then decreased to 18 °C and NEIL1 protein expression was induced by addition of 0.5 mM of isopropyl-D-1-thiogalactopyranoside (IPTG). After overnight growth, cells were harvested by centrifugation (6000 rpm, 4 °C, 20 min). The supernatant was discarded and cell pellets are stored at -20 °C until purified.

To purify NEIL1, cell pellets were resuspended in Lysis buffer (20 mM Tris-HCl pH 7.5, 500 mM NaCl, 5% w/v glycerol, 20 mM imidazole, 1% w/v NP-40, 0.1 mM PMSF, 5 mM beta mercapto-ethanol, protease inhibitor tab) using 5 mL of lysis buffer per gram of cell pellet. The cell suspension was homogenized and lysis was carried out by sonication at 4 °C (10 min, 5 seconds “on”, 10 seconds “off”, 50% power). The lysate was centrifuged at 50,000g for 45 min at 4 °C. The supernatant was collected and filtered at 0.45 um. This lysate was applied to a nickel affinity column (HisTrap FF 5mL ©Ge healthcare) pre-equilibrated with Ni-A buffer (20 mM Tris-HCl pH 7.5, 500 mM NaCl, 5% w/v glycerol, 20 mM Imidazole, 0.1 mM PMSF, 5 mM beta mercapto-ethanol). The resin was then washed with 20 CV of Ni-A buffer and the protein eluted with 10 CV of Ni-B buffer (Ni-A + 300 mM imidazole). Elution fractions were analyzed by SDS-PAGE and fractions containing NEIL1 protein were pooled together and diluted 2.5-fold with dilution buffer (20 mM HEPES pH 7.5, 5% w/v Glycerol, 1 mM EDTA, 0.1 mM PMSF, 2.5 mM beta mercapto-ethanol). The diluted protein solution was loaded onto a tandem of pre-equilibrated Q and S columns in buffer A (20 mM HEPES pH 7.5, 125 mM NaCl, 5% w/v glycerol, 1 mM EDTA, 0.1 mM PMSF, 2.5 mM beta mercapto-ethanol). At this ionic strength, the Q column filtered out binding contaminants while NEIL1 flows through the resin and then binds to the S column. Subsequently, a 20 CV wash step with buffer A was applied and the Q column was removed. The protein was eluted using a linear sodium chloride gradient from 200 mM to 1 M over 10 CV. Fractions were analyzed by SDS-PAGE, and those that contained pure NEIL1 were pooled together and concentrated using a Amicon © cutoff 10 kDa. The concentrated protein was applied to a S75 SEC column pre-equilibrated with S75 buffer (20 mM HEPES pH 7.5, 200 mM NaCl, 5% w/v glycerol, 1 mM EDTA, 0.1 mM PMSF, 2.5 mM Beta mercapto-ethanol). Fractions containing the final pure protein, as confirmed by SDS-PAGE, were pooled together, aliquoted, flash frozen in liquid nitrogen and stored at -80 °C.

### Nuclear Magnetic Resonance

All NMR experiments were recorded at 25 °C on Bruker Avance III NMR spectrometers operating at 600 or 800 MHz in a buffer containing 20 mM HEPES pH7.5, 200 mM NaCl, 5 mM DTT, and 5% D_2_O. Data were acquired from 200 μL samples in 3 mm NMR tubes. ^15^N-^1^H SOFAST HMQC spectra were recorded using 32 scans, 1024 points in the direct ^1^H dimension and 128 points in the indirect ^15^N dimension, with a recycle delay of 200 ms. All data were processed using Topspin© (Bruker) and analyzed using CCPNMR analysis software (20). Chemical Shift Perturbations (CSP) were calculated using:

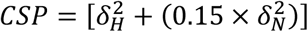

CSPs were plotted using pyplot.

### Multiple sequence alignment

Protein sequences were obtained from the uniprotKB portal (https://www.uniprot.org) (21) and aligned using Clustal OMEGA (https://www.ebi.ac.uk/Tools/msa/clustalo/). (22) The alignment figure was generated using JALVIEW. (23)

### Homology modelling of RPA32C in complex with NEIL1 RBS

The complex of RPA32C and the NEIL1 RPA Binding Motif (NEIL1 RBF) was generated using Modeller version 9.2 (24). The crystal structure of RPA32C with SMARCAL1 (PDB: 4MQV) was used as a template. 100 models were generated and the model with lowest DOPE score was kept as the final model.

### Isothermal titration calorimetry

Proteins samples were dialyzed against the same buffer solution containing 50 mM HEPES (pH 7.5), 150 mM NaCl, 10% glycerol and 0.5 mM TCEP. ITC data were collected using a Microcal VP isothermal titration calorimeter operating at 25 °C. Concentrations were 50 uM in the cell and 150 uM in the syringe, and the injection volume was 10 uL. Data were analyzed using NITPIC software, version 1.2.7 (25).

### Knockdown of NEIL1 using siRNA

U2OS (ATCC, HTB-96) and MCF7 cells (ATCC, HTB-22) were transfected using either an siRNA pool targeting NEIL1 (Dharmacon, L-008327-00-0005) or a non-targeting siRNA (Dharmacon, D-001810-01-20) as a control. These cells were transfected with siRNA using a Neon Transfection system (ThermoScientific, MPK5000) under conditions recommended by the manufacturer.

### FM-HCR analysis of DNA repair capacity in cells complemented with NEIL1

72-h after siRNA transfection, cells were transfected with 750 ng of either pMax_WT_NEIL1 or pMax_Mutant_NEIL1 using Lipofectamine3000 (ThermoScientific, L3000001). After an additional 24-h in culture, the cells were then trypsinized, collected by centrifugation, and transfected with Fluorescence Multiplex-Host Cell Reactivation (FM-HCR) reporter plasmids using the Gene Pulser MXCell Plate Electroporation System (Bio-Rad, #165-2670) using conditions 260 V, 950 μF. FM-HCR assays were carried out as previously described (26)). Reporter plasmids were prepared as a cocktail containing pmax_GFP plasmid damaged with 800 J/cm^2^ UVC radiation (herein referred to as GFP_UV) and an undamaged pMax_mPlum control as well as an undamaged cocktail containing pMax_GFP and pMax_mPlum. A control analysis was also completed using an mOrange-expressing plasmid containing a site-specific 8-oxoguanine lesion opposite a cytosine. The resulting fluorescence was then measured using an Attune NxT Flow Cytometer (ThermoScientific). NER capacity was also determined in HAP1 NEIL1 knockout cells obtained from (Horizon Discovery). These cells were transfected with 750 ng of an empty pMax vector (pMax_EV), pMax_WT_NEIL1, or pMax_Mutant_NEIL1 using Lipofectamine3000 and analyzed via FM-HCR as previously described (26).

### Clonogenic Survival Assay

NEIL1 KO cells were transfected with 750 ng pMax_EV, pMax_WT_NEIL1, or pMax_Mutant_NEIL1 using Lipofectamine3000 and allowed to incubate for 24-h. Cells were then collected and re-plated at 500 cells/well with either 5 μM 4-nitroquinolone *N*-oxide (4NQO) (Sigma, N8141) or a dimethyl sulfoxide (DMSO) (Sigma, D8418) control. The cells were left for 7 days and then fixed using 100% methanol (MeOH) (VWR, BDH2029) and stained with a crystal violet (Sigma, C6158) solution (20% MeOH, 80% H_2_O, 0.05% crystal violet). Visible colonies containing 50 or more cells were counted and recorded.

### In vitro assay for NEIL1 activity

To determine the functional activity of the WT and RPA-binding mutant NEIL1 variants, an *in vitro* enzyme activity assay was performed. Purified WT (0.1 μM) or RPA-binding mutant NEIL1 (0.1 μM) was incubated with 10 nM closed circular plasmid DNA encoding BFP which was previously irradiated with 100Gy X-ray radiation in a solution of 150 mM NaCl, 50 mM HEPES (4-(2-hydroxyethyl)-1-piperazineethanesulfonic acid) [pH 7.5], 1 mM DTT (Dithiothreitol), 5% (v/v) glycerol, 100 μM BSA (bovine serum albumin), and 1 mM EDTA (ethylenediaminetetraacetic acid). The reaction was allowed to proceed for 90 min at 37 °C. The negative control contained no purified enzyme to ensure any observed digestion was, in fact, due to the presence of NEIL1 activity. Each reaction was immediately run on a 1.0% agarose gel (80 min, 125 V) and imaged using an iBright FL1500 Imaging System.

## Results

### RPA interaction with NEIL1 is mediated by RPA 32C and 70AB domains

To identify which domains of RPA interact with NEIL1, two-dimensional ^15^N-^1^H Heteronuclear Single Quantum Coherence (HSQC) NMR experiments were acquired to monitor perturbations of backbone ^15^N and ^1^H chemical shifts (CSPs). Following the strategy of past studies characterizing RPA interaction partners, ^15^N-enriched samples were prepared for the two protein recruitment domains RPA 32C and 70N, and the tandem high affinity DNA binding domains RPA70AB. In these experiments, the binding of 44 kDa NEIL1 is expected to result in substantial line broadening of the signals from the RPA domain(s).

Previous studies mapped binding of NEIL1 to the RPA70 subunit, which contains the RPA70N protein recruitment domain (12). However, upon addition of up to 4 molar equivalents of NEIL1, no significant perturbations of the spectrum of RPA70N were observed **(Figure 1 A)**. In contrast, addition of NEIL1 to ^15^N-enriched RPA 32C resulted in substantial line broadening throughout the whole spectrum **(Figure 1 A)**, indicating a significant interaction. To obtain deeper insights, binding of RPA32C by NEIL1 was characterized by Isothermal Titration Calorimetry (ITC) **(Figure 1 B)**. These data revealed that binding was a primarily endothermic enthalphy-driven process with a dissociation constant (Kd) of 240 nM.

**Figure 1:**
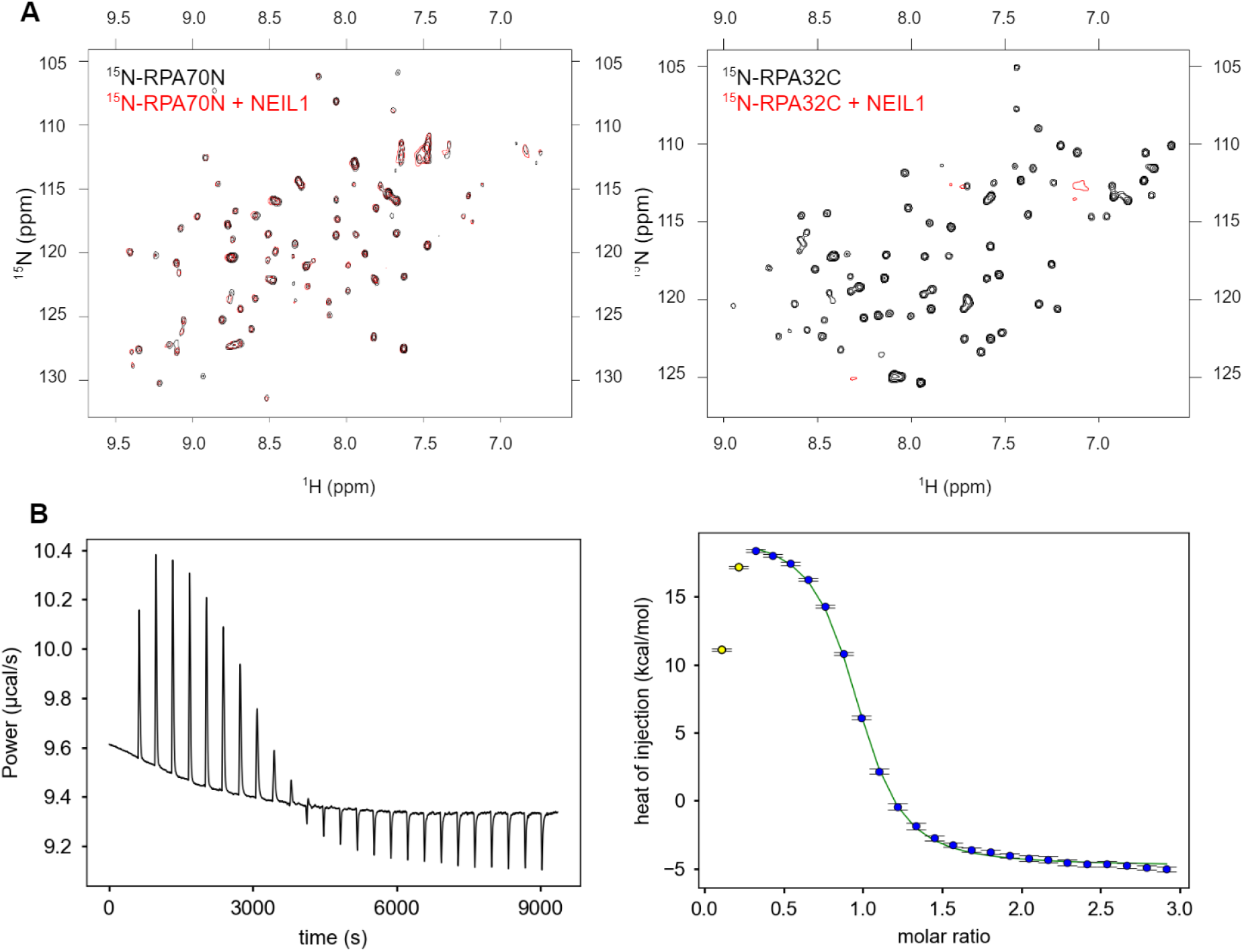
Interaction of NEIL1 with the RPA70N and RPA32C protein recruitment domains. A-^15^N-^1^H HSQC spectra recorded at 800 MHz of ^15^N-enriched RPA70N (left) and RPA32C (right) in the absence (black) or presence (red) of 4 molar equivalents of NEIL1. Spectra were acquired at 25 °C in a buffer containing 25 mM Tris-HCl pH 7.5, 150 mM NaCl, and 1 mM DTT. B-Isothermal titration calorimetry experiment for RPA32C titrated with NEIL1. Data were acquired at 25 °C in a buffer containing 50 mM HEPES pH7.5, 150 mM NaCl, 10% glycerol, and 0.5 mM TCEP. A Kd value of 240 nM was extracted from the data for RPA32C.

In previous studies we often observed that the interaction of RPA with partner proteins have, in addition to a readily characterized interaction with RPA70N or RPA32C, a secondary interaction with the tandem high affinity DNA binding domains RPA70AB (e.g. (27)). These interactions are invariably weaker than the contacts with the corresponding protein recruitment domain. An interaction of NEIL1 with RPA70AB would also support the previous report of NEIL1 interacting with the RPA70 subunit (12). To test this hypothesis, the ^15^N-^1^H HSQC spectrum of ^15^N-enriched RPA70AB was recorded with increasing amounts of NEIL1 **(Figure 2)**. This titration produced a gradual disappearance of RPA70AB signals due to line broadening. The titration was continued to an 8-fold excess of NEIL1, yet spectral changes continued to occur. The inability to saturate the effect indicates the interaction of NEIL1 with RPA70AB is weaker than the interaction with the RPA32C domain. Attempts to characterize the thermodynamics of binding by ITC revealed very small heats of binding and far from complete titration even at 2.5-fold molar excess, consistent with the weak binding affinity inferred from the NMR experiments.

**Figure 2:**
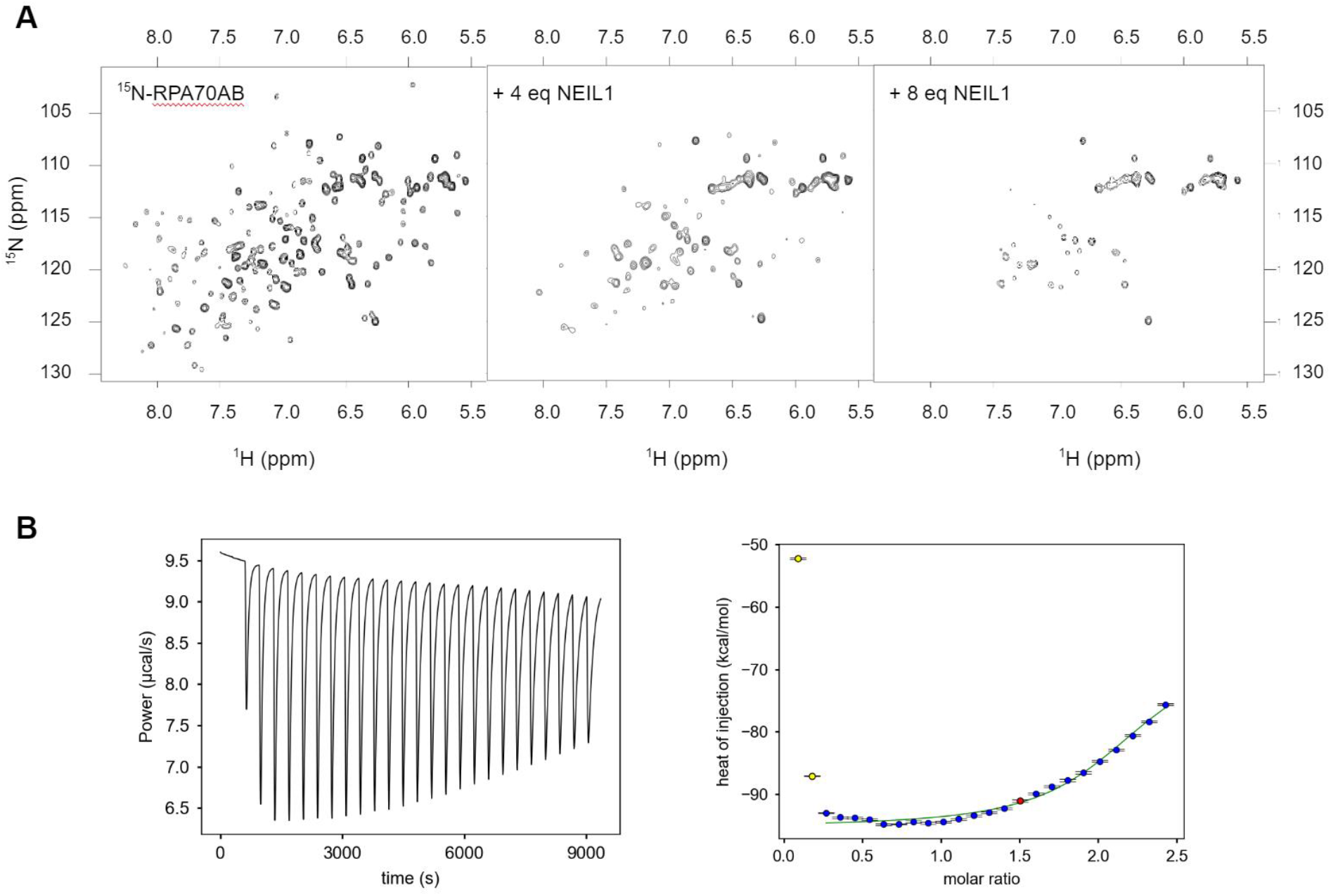
Interaction of NEIL1 with the tandem high affinity DNA binding domains RPA70AB. A-^15^N-^1^H HSQC spectra recorded at 800 MHz of ^15^N-enriched RPA70AB free and in the presence of 4 and 8 equivalents of NEIL1. Spectra were acquired at 25 °C in a buffer containing 25 mM Tris/HCl pH 7.5, 150 mM NaCl, and 1 mM DTT. B-Isothermal titration calorimetry experiment for RPA70AB titrated with NEIL1. Data were acquired at 25 °C in a buffer containing 50 mM HEPES pH7.5, 150 mM NaCl, 10% glycerol, and 0.5 mM TCEP. The binding of RPA70AB is so weak that it is not possible to quantify the thermodynamics of binding.

### The high affinity RPA binding motif is contained in the NEIL1 Common Interaction Domain

All NEIL1 interactions with proteins characterized to date occurs through the disordered C-terminal region (NEIL1_290-390_) termed the Common Interaction Domain (CID). In order to locate more precisely the high affinity RPA binding motif of NEIL1, a multiple sequence alignment was generated of NEIL1 CID with known RPA 32C binding motifs **(Figure 3)**. This allowed us to identify a putative RPA-binding motif in the first half of the NEIL1 CID, within NEIL1 residues 305 to 331.

**Figure 3:**
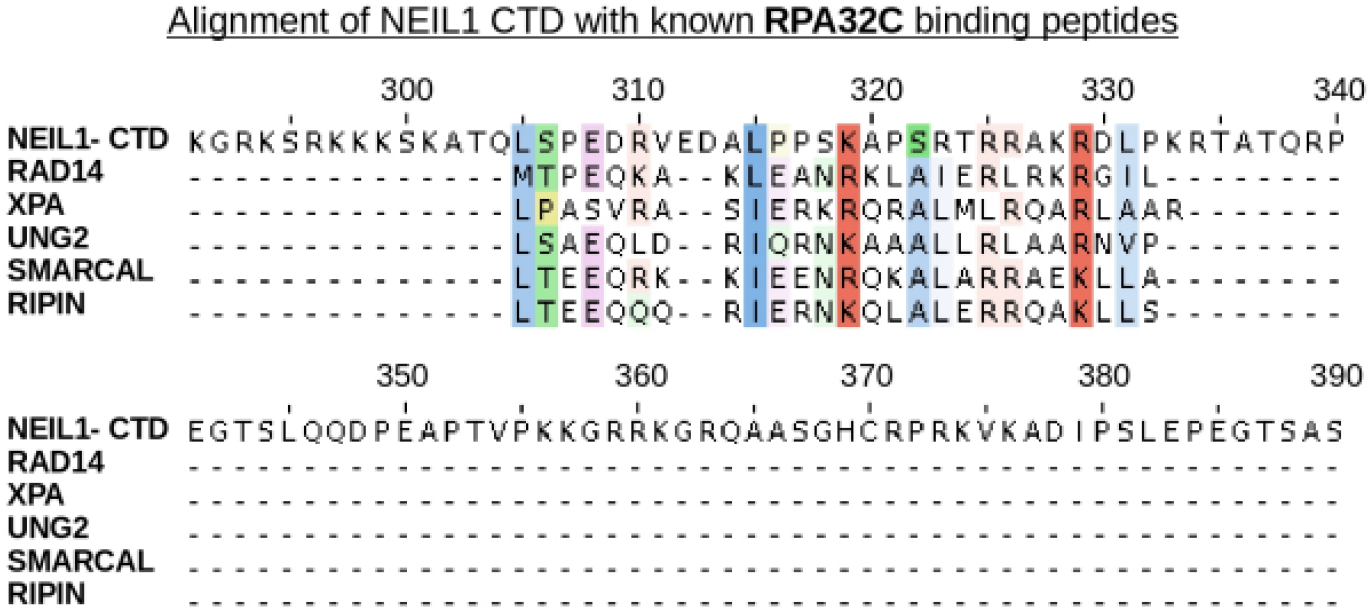
Primary sequence alignment of NEIL1 CTD with known RPA32C binding motifs. The NEIL1 C-terminal domain (residues 290-390) was aligned with known RPA32C binding motifs from various known RPA32C partner proteins. The degree of transparency of each color is proportional to the extent of conservation. Hydrophobic residues are colored in blue, positively charged residues in orange, negatively charged in pink, and polar residues in green.

To test this hypothesis, ^15^N-^1^H HSQC NMR spectra of ^15^N-enriched RPA32C were acquired in absence or presence of a fragment of NEIL1 CID containing the putative RPA binding motif (NEIL1_289-349_). Comparison of the two spectra revealed significant CSPs throughout the RPA32C spectrum **(Figure 4AB)**, indicating that NEIL1_289-349_ does indeed contain the RPA interaction motif. Mapping of the CSPs onto the structure of RPA32C shows that the most strongly perturbed signals correspond to residues located at the canonical interface between RPA32C and known binding partners **(Figure 4C)**. Based on these results, it was evident that a high-quality homology model of the complex of RPA32C and the NEIL1 RPA binding motif could be generated using the co-crystal structure of the complex of RPA32C with a peptide fragment of SMARCAL1 (PDB: 4MQV) as a template.

**Figure 4:**
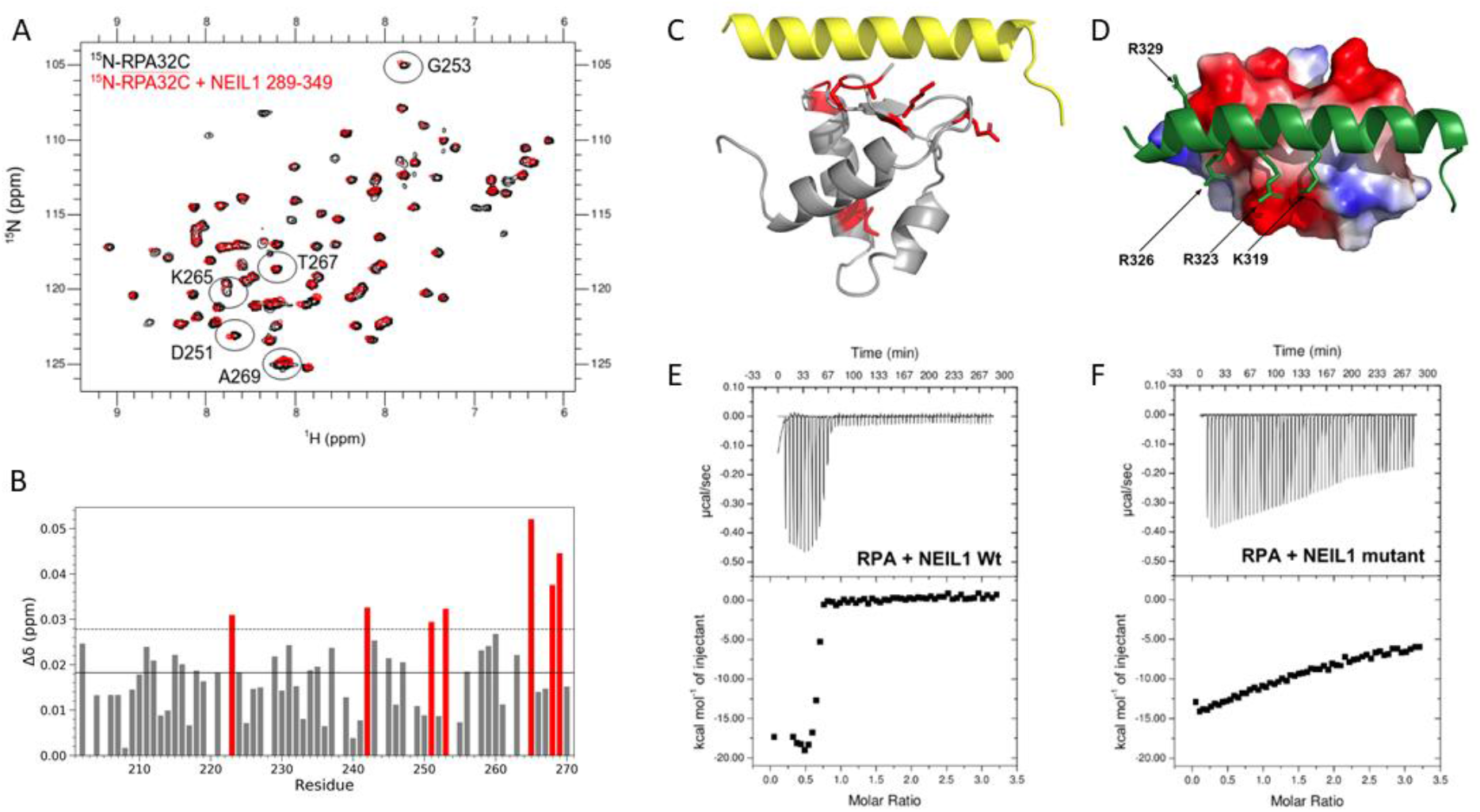
Mapping and Disruption of NEIL1 Interaction with RPA32C. A-^15^N-^1^H HSQC spectra recorded at 800 MHz of ^15^N-enriched RPA32C in the absence (black) or presence (red) of 4 molar equivalents of NEIL1_289-349_. Spectra were acquired at 25 °C in a buffer containing 25 mM Tris-HCl pH 7.5, 150 mM NaCl, and 1 mM DTT. The resonances with the largest CSPs are identified by circles and residue number. <u>B-</u> Plot of NMR CSPs as a function of residue number. The mean of all CSPs is identified by the solid line and the standard deviation over the mean is identified by the thin dotted line. The most significantly perturbed residues are colored red. C-Map of the CSPs on the homology model of the complex of RPA32C and the NEIL1_289-349_. The same significantly perturbed residues as in B are colored red in the model, showing they map primarily to the known RPA32C interaction interface. D-Electrostatic field of RPA32C shown on the homology model of the complex with NEIL1_289-349_ (green). Surfaces are colored red for negative charge, white for neutral and blue for positive charge. The basic side chains in NEIL1 that complement the acidic RPA32C surface and were mutated are identified with arrows. E-F-Isothermal titration calorimetry of RPA titrated with NEIL1 wild-type (panel E) and mutant (panel F). The data were acquired at 25 °C in a buffer containing 50 mM HEPES (pH 7.5), 150 mM NaCl, 10% glycerol and 0.5 mM TCEP. The dissociation constant (Kd) for wild-type NEIL1 is 20 nM, which is consistent with a previously reported value (34). The binding of the mutant is so weak that the binding affinity cannot be determined by this approach.

### Structure based mutations of NEIL1 CID specifically disrupt the interaction

The homology model of the RPA32C-NEIL1_289-349_ complex was then used to examine the interface **(Figure 4C)**. A combination of both electrostatic and hydrophobic interactions are found **(Figure 4D)**. The electrostatic component involves the acidic surface of RPA32C complemented by multiple basic residues at the surface of NEIL1. Since electrostatic interactions are long-range in nature and the interface between the two proteins covers a significant surface area, a multi-site charge reversal mutation was designed, which we surmised would specifically inhibit the interaction between NEIL1 and RPA. To test this hypothesis, mutations K319E, R323E, R326E and R329E were introduced into NEIL1. An ITC titration was then performed to compare the interaction of the wild-type and mutant NEIL1 proteins with RPA. Comparison of the two titrations revealed that the mutations in NEIL1 severely impede its interaction with RPA **(Figure 4EF)**.

### Loss of NEIL1-RPA Interaction Results in a Mild Nucleotide Excision Repair Defect

To assess the functional relevance of the RPA-NEIL1 interaction, a targeted knockdown of NEIL1 was carried out using both U2OS and MCF7 cells. 96-h post-transfection with either non-targeting siRNA or siRNA targeting NEIL1, cells were collected and transfected using FM-HCR reporter plasmids. After 24-h, the cells were analyzed via flow cytometry. This analysis revealed an unexpected increase in repair of 8oxoG:C lesions that are excised by NEIL1, albeit less efficiently than by OGG1 (**Figure 6A**). Our data suggest that although NEIL1 can stimulate turnover of OGG1 after the base excision step,(28) slower excision of 8oxoG:C lesions by NEIL1 can compete with more rapid initiation by OGG1, resulting in less efficient initiation of BER. Surprisingly, our analysis also revealed a significant decrease in NER capacity in cells depleted for NEIL1 (**Figure 6B**).

To identify the potential importance of the interaction of NEIL1 and RPA, cells transfected with the non-targeting siRNA or NEIL1-targeting siRNA were also transfected with either pMax_WT_NEIL1 or pMax_Mutant_NEIL1 72-h after the initial siRNA transfection and analyzed for repair capacity using FM-HCR. Comparison of cells transfected with pMax_WT_NEIL1 and pMax_Mutant_NEIL1 showed that complementation with the WT protein was able to rescue the higher NER capacity and lower excision of 8oxoG:C, whereas the mutant NEIL that lacks RPA-binding capability rescued repair of the 8oxoG:C lesion (**Figure 5A**), but not the NER defect (**Figure 5B**). An *in vitro* plasmid-based biochemical assay with purified proteins confirmed that both WT and mutant NEIL1 are catalytically active against oxidative lesions induced by ionizing radiation (**Figure 5C**). Both variants of the bifunctional glycosylase converted closed circular plasmid with oxidative damage induced by ionizing radiation (Lane 2) into open circular DNA with efficiency similar to each other (Lanes 3,4). Taken together, these data suggest that RPA binding is dispensable for NEIL1-dependent recognition and processing of 8oxoG:C *in vivo* and excision of radiation-induced oxidative lesions *in vitro*, but is required for an apparent role for NEIL1 in the NER pathway.

**Figure 5:**
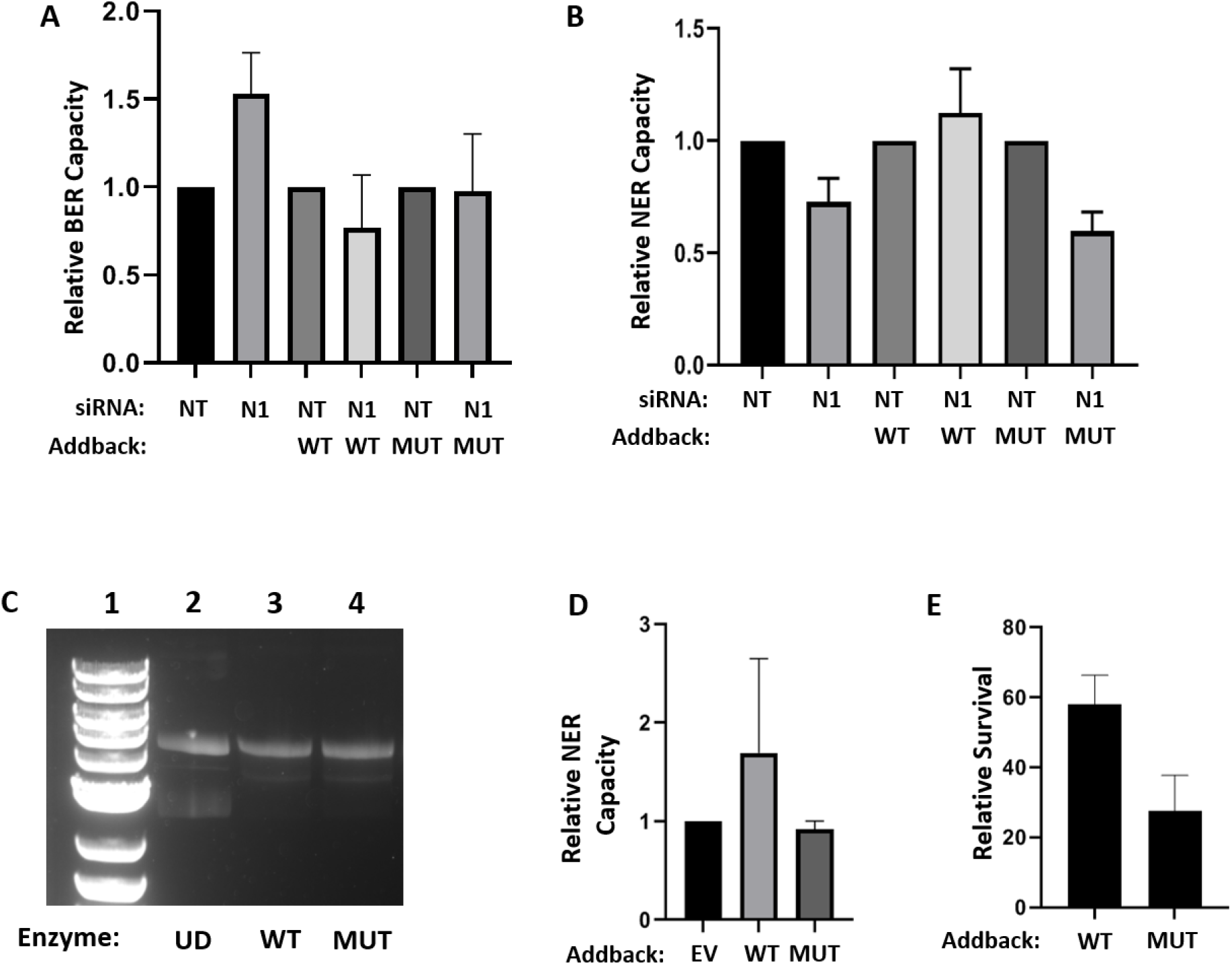
DNA repair capacity and sensitivity to 4NQO. A. FM-HCR analysis of 8oxoG:C base excision in U2OS cells complemented with wild type (WT) or mutant (MUT) NEIL1 following siRNA-mediated depletion of NEIL1. B. FM-HCR analysis of NER in U2OS cells complemented with wild type (WT) or mutant (MUT) NEIL1 following treatment with non-targeting siRNA (NT) or siRNA-mediated depletion of NEIL1 (N1). C. Gel electrophoretic analysis of a plasmid treated with 1000 Gy ionizing radiation (Lane 2) following treatment with WT or MUT NEIL1 (Lanes 3 and 4). D. FM-HCR analysis of NER capacity in HAP1 NEIL1 KO cells transfected with an empty vector (EV) control or complemented with WT or MUT NEIL1. E. Clonogenic survival assay of HAP1 NEIL1 KO cells complemented with WT or MUT NEIL1 and treated with 5 nM 4NQO.

To reproduce these observations in a model that does not rely on siRNA-mediated depletion of NEIL1, we used HAP1 cells in which NEIL1 has been deleted using CRISPR-Cas9. These KO cells were transfected using either pMax_EV, pMax_WT_NEIL1 or pMax_Mutant_NEIL1. Consistent with the results in U2OS and MCF7, complementation with the WT NEIL1 that is able to bind RPA results in a 2-fold increase in NER capacity, whereas complementation with the empty vector or the RPA-binding-deficient NEIL1 mutant did not (**Figure 5D**).

Finally, to confirm that the modest changes in NER observed using plasmid-based assays are biologically relevant, we used a clonogenic survival assay following exposure of cells to 4NQO, which induces bulky DNA lesions that are repaired by the NER pathway. NEIL1 KO cells complemented with pMax_WT_NEIL1 were significantly more resistant to 4NQO than NEIL1 KO cells complemented with pMax_Mutant_NEIL1. By contrast, WT control cells transfected with either plasmid showed similar sensitivity. (**Figure 5E**). These data suggest that the NEIL1-RPA interaction protects cells from killing by 4NQO by promoting NER of the bulky DNA adducts produced in genomic DNA.

## Discussion

Since the discovery that NEIL1 is up-regulated in S phase and is also able to process ssDNA (10, 29, 30), there has been a great deal of interest in understanding its role in replication associated DNA repair (11, 31–33). As the BER pathway requires generation of potentially toxic DNA structures in the context of replication, such as abasic sites and strand breaks, it is essential that these two cellular processes are regulated and coordinated. The coordination between the two DNA processing pathways relies on a network of key protein-protein interactions, such as that between RPA and NEIL1 as characterized here. Understanding how these interactions occur at the molecular level provides valuable insights for testing functional models and elucidating the mechanism of the interplay between replication and BER.

In this study, we have shown that unlike inferences drawn in a previous study (34), the primary contact between NEIL1 and RPA is the RPA32C protein recruitment domain. We have also confirmed the existence of an interaction with the RPA70 subunit as reported previously (35), which we mapped to the tandem RPA70AB domains and showed by ITC that it is considerably weaker than the interaction with RPA32C. A homology model of the complex of RPA32C and the NEIL1 RPA binding motif in the CID was then used to design specific mutations at the NEIL1-RPA interface to inhibit the interaction. We went on to show that a NEIL1 variant with four charge-reversal mutations effectively suppresses the physical interaction with RPA. This NEIL1 mutant serves as a valuable reagent for investigating the functional significance of the NEIL1 interaction with RPA and its effect on the coupling of replication and BER, both in vitro and in cells.

The relevance of the interaction between RPA and NEIL1 is underscored by the recent discovery that in mitochondria the functional homolog of RPA, mtSSB, also interacts with NEIL1 (36). Moreover, mtSSB interacts with NEIL1 in the same region of the NEIL1 CID as RPA. Here we have shown that the molecular basis of the interaction between NEIL1 and RPA32C is very similar to the previously characterized interaction of another DNA glycosylase involved in the initiation of the BER pathway, UNG2 (35). Like the NEIL1 CID, UNG2 has an RPA32C binding motif in a disordered domain adjacent to the catalytic domain, implying there may be a similar mechanism of binding. A recent study using IPOND (isolation of protein on nascent DNA) revealed that NEIL1, UNG2 and other glycosylases are found at replication forks (32). Different glycosylases function to repair different types of DNA base damage. Hence, the ready availability of glycosylases at the replication fork and the need to rapidly repair damaged bases without accumulating repair intermediates that are more toxic than the initial base lesions may be regulated by their interaction with RPA. RPA may stimulate pre- and post-replicative BER in dsDNA flanking ssDNA but inhibit BER in ssDNA, preventing formation of toxic strand break.

Finally, it appears that although the interaction between NEIL1 and RPA is dispensable for the activity of NEIL1 against oxidative lesions in duplex DNA, the interaction plays an unexpected role in cellular NER capacity. Knockdown of NEIL1 results in a mild NER defect that is only rescued when NEIL capable of binding RPA is re-introduced to these cells. This is supported through the introduction of these same proteins into NEIL1 KO cells using both FM-HCR and a clonogenic survival assay measuring sensitivity to 4NQO, which induces bulky DNA adducts that are repaired by the NER pathway. Collectively, the data are consistent with rescue of NER proficiency only when NEIL1 is capable of binding of RPA. Our results add to previous work indicating crosstalk between DNA glycosylases and NER (37–39). Noting that NEIL1 is capable of binding some types of bulky DNA adducts(39), our data may suggest a possible handoff mechanism wherein NEIL1 recruits XPA to sites of DNA damage via shared interactions with RPA. In conclusion, our findings provide new insights into the role of interactions between RPA and NEIL1 in multiple DNA repair pathways, and suggest previously unrecognized targets for therapeutic inhibition of DNA repair in cancers.

## Acknowledgement

We would like to thank Drs. Muralidhar Hegde and Sankar Mitra for providing NEIL1 expression vectors and extensive background, as well as advice, stimulating discussions and motivation to pursue this project. This work was supported by the US National Institutes of Health (R35 GM118089 and R01 CA092584 to WJC, P01 CA092584 to ZDN and WJC; U01 ES029520 and P30 ES000002 to ZDN). The NMR facilities were supported by NIH grant (S10 RR025677).

